# Separating faces in ARMS metabarcoding improves marine biodiversity monitoring: a comparison across protocols, experimental designs, and photographic surveys

**DOI:** 10.1101/2025.11.24.690150

**Authors:** Anne Chenuil, Elyna Bouchereau, Térence Legrand, Virgile Calvert, Cécile Chemin, Sandrine Chenesseau, Dorian Guillemain, José Miguel Gutiérrez Ortega, Anne Haguenauer, Michèle Leduc, Florent Marschal, Christian Marschal, Fatma Mirleau, Marjorie Selva, Laurent Vanbostal, Frédéric Zuberer, Pascal Mirleau, Laetitia Plaisance, Vincent Rossi, Sandrine Ruitton, Emese Meglécz, Vincent Dubut

## Abstract

Monitoring marine biodiversity requires approaches that capture its full complexity through space and time. DNA metabarcoding coupled with Autonomous Reef Monitoring Structures (ARMS) is increasingly used for this purpose, yet most applications still pool all sessile fractions and rarely benchmark molecular ouputs against photographic observations. Here, we combined photographic analysis with cytochrome c oxidase I (COI) metabarcoding across ten north-western Mediterranean sites to test, compare, and refine ARMS-based monitoring protocols. We first optimized laboratory procedures (DNA extraction and polymerase choice) and applied the control-driven, replicate-aware VTAM pipeline to minimize false positives and ensure full traceability. We then conducted the first face-by-face comparison of α- and β-diversity between imaging and eDNA in which each individual ARMS face was metabarcoded separately rather than pooled. Metabarcoding detected ∼15× higher site-level richness and revealed stronger correlations with geographic distance and environmental gradients, whereas photography provided complementary information on macro-taxa and surface cover. For metabarcoding, processing each face separately yielded much higher richness and markedly stronger β-diversity–distance correlations than with the NOAA pooling protocol, demonstrating that pooling inflates sampling variance resulting in a loss of the ecological signal. Grouping faces into five structural categories offered a more operational alternative while further increasing α-diversity and strengthening β-diversity correlations. Overall, our results show that retaining ARMS microhabitat structure is critical for maximizing metabarcoding performance. Using five structural sessile fractions per ARMS combined with a control-driven bioinformatic workflow provides a reproducible, scalable framework for long-term eDNA monitoring and early detection of biodiversity change.

## 1. Introduction

Biodiversity protection is increasingly recognized as a global priority (Cowie et al., 2022). Global assessments warn that current biodiversity loss is unprecedented and threatens vital ecosystem services (Díaz et al., 2019a, 2019b). Marine ecosystems face intensifying pressures from climate change, ocean acidification, deoxygenation, overfishing, pollution, and habitat destruction (Halpern et al., 2015; Duarte et al., 2020). Effective protection therefore depends on timely, evidence-based monitoring of species composition and ecological change (Estes et al. 2021; Miloslavich et al., 2018).

Meeting this need requires standardized, sensitive, and scalable methods that capture taxonomic and genetic diversity and they associated spatio-temporal dynamics across broad geographic ranges (Laikre et al., 2020). For hard-bottom benthic communities, two major challenges remain: (i) developing standardized sampling schemes and (ii) ensuring consistent, taxonomically reliable assessments (Danovaro et al., 2016, 2017). Historically, benthic assemblages were identified morphologically by taxonomic experts, which is a labor-intensive process that depends on a diminishing pool of experts (Appeltans et al., 2012; Costello et al., 2013) and is prone to inconsistencies among experts, limiting reproducibility (Aylagas et al., 2016). These limitations hinder the long-term and large-scale monitoring plans required to track biodiversity change. The challenge is particularly acute in regions of exceptionally high species richness, such as the Mediterranean and many tropical seas, where small, morphologically similar or cryptic taxa dominate hard-bottom assemblages (Coll et al., 2010; Bianchi et al., 2022; Obura, 2012). In these biodiversity hotspots, the sheer number of species and the prevalence of cryptic or early-life stages make comprehensive morphological inventories impractical.

Autonomous Reef Monitoring Structures (ARMS) offer a promising solution for standardized hard-bottom sampling. Initially developed in tropical seas (Plaisance et al., 2011), they are now also deployed in temperate and polar environments (Couëdel et al., 2023; David et al., 2019; Obst et al., 2020; Pearman et al., 2018; Daraghmeh et al., 2025; Pagnier et al., 2025). ARMS provide a uniform, replicated habitat that becomes colonized by diverse sessile and mobile organisms. Their community composition has typically been characterized either through photography-based analyses of the 17 exposed plate faces (David et al., 2019) or through DNA metabarcoding of the organisms scraped from the structure after retrieval (Leray & Knowlton, 2015), and from their mobile fractions. The global ARMS program details a step-by-step protocol from deployment to data analysis, ensuring a globally standardized methodology. Therefore, ARMS deployment is suitable for large-scale network initiatives such as the Marine Biodiversity Observation Network (ARMS-MBON; Obst et al., 2020; Pearman et al., 2020) that integrates photography analyses and metabarcoding across European seas and polar regions, providing long-term, FAIR biodiversity data and delivering Essential Biodiversity Variables (EBVs; Miloslavich et al., 2018) as well as early detection of non-indigenous species.

Both photography-based and metabarcoding approaches have advantages for biodiversity assessment and can be partly automated thanks to advances in artificial intelligence and next-generation sequencing, respectively (Beijbom et al., 2015; Pearman et al., 2018; Deiner et al., 2017; Taberlet et al., 2018). Furthermore, beyond community inventories, ARMS metabarcoding can also capture intraspecific genetic diversity and reveal cryptic population structure (Thomasdotter et al., 2023). On one hand, metabarcoding typically identifies taxa from bulk DNA extracts using universal primers and high-throughput sequencing (e.g. Andújar et al., 2018; Brandt et al., 2021a; Pearman et al., 2020). However, false positives and false negatives can be generated without careful quality control (Zinger et al., 2019; Alberdi et al., 2018; Brandt et al., 2021b). Here we apply a stringent, control-driven pipeline (VTAM; González et al., 2023) that minimizes such errors and ensures full traceability. On the other hand, photography-based analyses can reach species-level resolution, for some taxa, when performed by trained experts, but large monitoring networks often require coarser, higher-taxonomic classifications (David et al., 2019; Pearman et al., 2018; Miloslavich et al., 2018).

The standard NOAA protocol for metabarcoding uses the two smallest mobile fractions (100–500 µm and 500–2000 µm) separately and a single composite sessile fraction scraped from all nine ARMS plates (Leray & Knowlton, 2015; Pearman et al., 2020), thereby sacrificing microhabitat-scale heterogeneity among individual plate faces (David et al. 2019). To overcome this limitation and enhance ARMS-based biodiversity assessments, we refined existing protocols through three key methodological improvements. First, we evaluate laboratory procedures by comparing two DNA extraction kits and two polymerases to identify the most efficient combination for diverse marine assemblages. Second, we implement a face-by-face metabarcoding design in which the sessile community of each of the 17 ARMS faces is analyzed individually rather than as a single pooled sample. This enables the first direct face-by-face comparison of hard-bottom community composition using both photography-based annotation and COI metabarcoding, providing unprecedented cross-validation of α- and β-diversity estimates. Third, we use two additional ARMS for each site where metabarcoding used pooled sessile fractions, to test whether retaining face-level resolution improves richness detection and strengthens correlations of community dissimilarity with environmental variables (geographic distance and environmental gradients relative to the standard pooled NOAA protocol). Finally, we tested a more operational scheme in which five sessile fractions per ARMS (corresponding to distinct microhabitats) were analyzed separately while including three ARMS units per site.

Here, we evaluate whether separating ARMS faces for metabarcoding improves biodiversity assessments relative to pooled designs. Specifically, we compare four monitoring protocols—one based on photography (images taken on the 17 individual plate faces of each ARMS) and three based on metabarcoding (full face-by-face processing of the same 17 faces, pooling all faces per ARMS, and grouping faces into five structural categories)—to quantify their ability to recover α-, β-, and γ-diversity and to detect geographic and environmental structuring of communities. Because ARMS provide contrasting microhabitats (upward-vs. downward-facing surfaces, open vs closed compartments, varying exposure to light), we hypothesized that retaining face-level resolution would enhance the ecological signal recovered by metabarcoding. Accordingly, we predicted that: (i) metabarcoding of separated faces would reduce the impact of sampling variance and detect higher richness than pooled protocols, and (ii) geographic and environmental gradients would be more strongly recovered when ARMS faces are analyzed individually or by structural category.

## 2. Material and Methods

### 2.1. Field sampling and ARMS sampling design

Autonomous Reef Monitoring Structures (ARMS) were deployed by scuba divers at 12 sites distributed across three regions of the French Mediterranean coast (Figure 1) at depths of 16–22 m (Table 1). Units were immersed between 27 February and 7 April 2017 and retrieved between 5 March and 9 May 2018 (Table 1). Before dismantling and photographic documentation, each ARMS was kept in aerated seawater for 1 h to overnight. To remove vagile fauna, each plate was gently shaken prior to photography. After imaging, the sessile benthos was scraped from each face, preserved in 96 % ethanol, and stored in at +4°C until DNA extraction.

**Figure 1:**
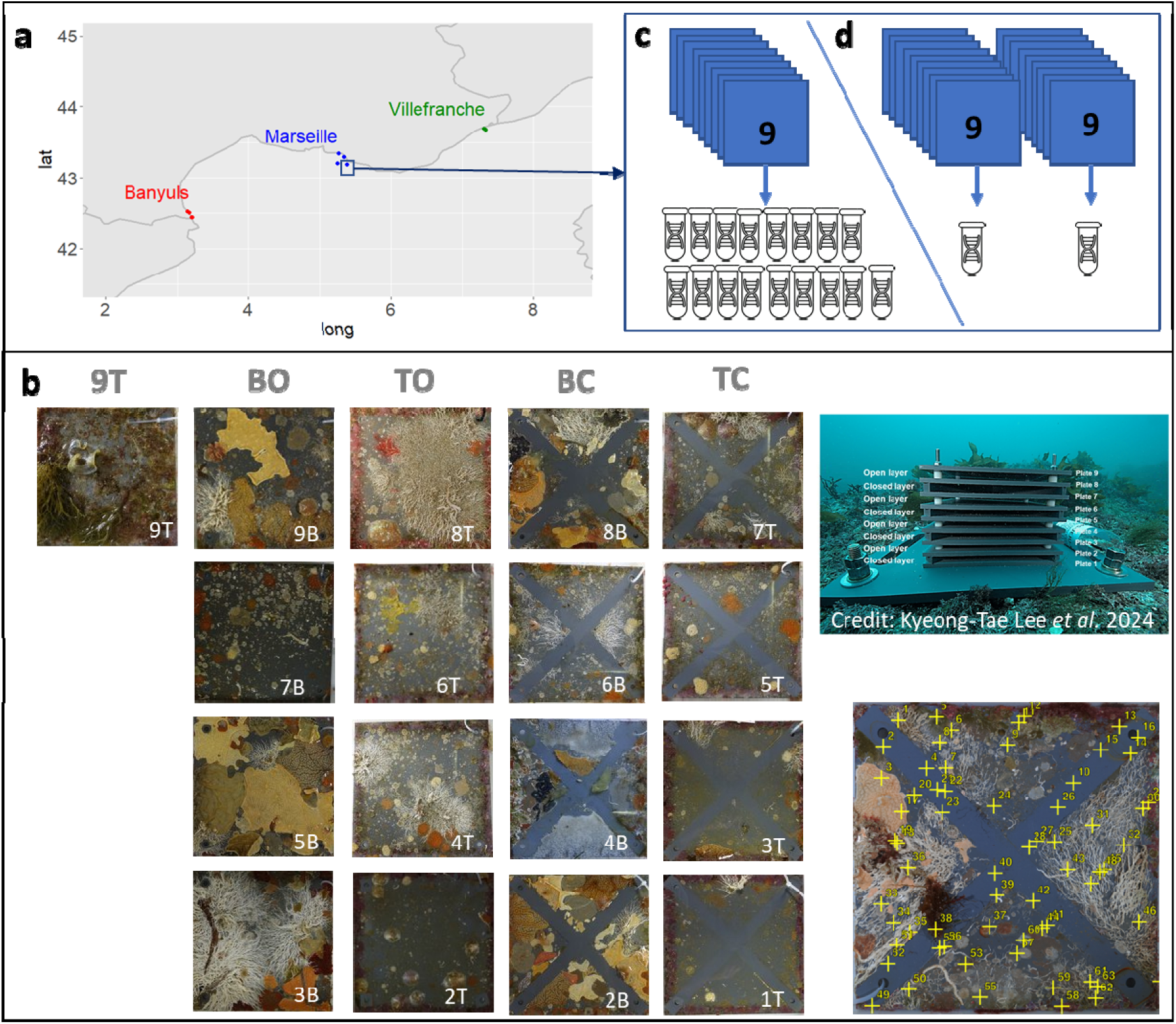
Study sites and experimental design for the 2018 dataset. The survey covered 12 sites distributed across three regions of the north-western Mediterranean coast (France). Each site hosted three Autonomous Reef Monitoring Structures (ARMS), and each ARMS comprised nine plates providing 17 colonizable faces. (a) Map of the 12 study sites grouped by region. (b) Photography-based analysis: images of the 17 faces from a single ARMS, arranged in five columns representing the main structural face categories (9T, BO = bottom open, TO = top open, BC = bottom closed, TC = top closed). Bottom right, an example of random points overlaid on a photograph for taxonomic labelling by an expert assisted by artificial intelligence. (c–d) Metabarcoding design for one of the ten sites where one ARMS was analyzed face by face. (c) 17 separate sessile samples corresponding to the 17 faces, allowing direct comparison with the photographic data. (d) Pooled sessile fractions from each of the other two ARMS at the same site, processed as a single metabarcoding sample following the standard NOAA protocol. Only 10 sites followed the design illustrated in this figure and in Table 1 were used for comparative purposes.

**Table 1:** Study sites and sampling details. For each site, the table reports depth, geographical coordinates (decimal latitude and longitude), dates of immersion and emersion, and the number of ARMS processed either as individual plate faces or as pooled sessile fractions for metabarcoding (numbers of samples are given in parentheses when duplicate analyses were performed on a single ARMS). Note: Although most analyses in this study include only 10 sites (where 17 separate sessile community metabarcoding were carried out for one ARMS), we present all 12 sites for overall information on the SEAMoBB project.

| Site code | Site name | Region | Latitude | Longitude | Depth | Date in | Date out | Pooled ARMS | 17 faces ARMS |
| --- | --- | --- | --- | --- | --- | --- | --- | --- | --- |
| BAN1 | Port-vendres S | Banyuls-sur-Mer | 42,51327 | 3,13743 | 17 m | 27/02/2017 | 17/04/2018 | 3 | 0 |
| BAN2 | Port-vendres N | Banyuls-sur-Mer | 42,52518 | 3,11050 | 17 m | 28/02/2017 | 12/04/2018 | 2 | 1 |
| BAN3 | Cerbère N | Banyuls-sur-Mer | 42,44818 | 3,17212 | 17 m | 28/02/2017 | 19/03/2018 | 2 | 1 |
| BAN4 | Cerbère S | Banyuls-sur-Mer | 42,44262 | 3,17405 | 17 m | 01/03/2017 | 19/03/2018 | 2 | 1 |
| MAR1 | Veyron | Marseille | 43,20683 | 5,25350 | 17,5 m | 16/03/2017 | 07/05/2018 | 2 | 1 |
| MAR2 | Digue des catalans | Marseille | 43,29167 | 5,34558 | 19 m | 20/03/2017 | 05/03/2018 | 2 | 1 |
| MAR3 | Niolon | Marseille | 43,34150 | 5,26525 | 16,5 m | 03/04/2017 | 09/05/2018 | 2 (3) | 1 |
| MAR4 | Plane | Marseille | 43,19067 | 5,38233 | 17,5 m | 07/04/2017 | 05/03/2018 | 2 (3) | 1 |
| VIL1 | Pointe de la gavinette | Villefranche-sur-Mer | 43,68683 | 7,32138 | 20 m | 27/03/2017 | 13/03/2018 | 2 | 1 |
| VIL2 | Phare du cap ferrat | Villefranche-sur-Mer | 43,67427 | 7,32803 | 18 m | 28/03/2017 | 15/03/2018 | 2 | 1 |
| VIL3 | Grotte à corail | Villefranche-sur-Mer | 43,68945 | 7,30883 | 17 m | 28/03/2017 | 13/03/2018 | 3 | 0 |
| VIL4 | Cap ferrat est | Villefranche-sur-Mer | 43,67675 | 7,33508 | 18 m | 29/03/2017 | 14/03/2018 | 2 | 1 |

Each ARMS consisted of nine superimposed plates (Figure 1), with an upper face (oriented upward) and a lower face (oriented toward the seafloor). In alternating inter-plate spaces, crossbars subdivided the gap into four compartments (see Pearman et al., 2020). Plate faces were designated by plate number and orientation (e.g. 1T, 2B, 2T … 9T, where “T” denotes the upward-facing surface and “B” the downward-facing surface). For subsequent statistical analyses, we grouped the 17 faces into five structural categories:

- 9T, the uppermost face;
- BC (bottom closed): faces 2B, 4B, 6B, 8B;
- BO (bottom open): faces 3B, 5B, 7B, 9B;
- TC (top closed): faces 1T, 3T, 5T, 7T;
- TO (top open): faces 2T, 4T, 6T, 8T.

This hierarchical face-type classification was used to test how fine-scale microhabitat influences community composition, and to evaluate whether grouping faces by structural categories before DNA extraction is an efficient approach for metabarcoding-based monitoring.

To assess the benefits of face-by-face sampling and compare imaging with metabarcoding at the finest level, we analyzed at each site: (i) one ARMS unit with all individual faces processed separately, and (ii) two ARMS units with all sessile material pooled into a single DNA extraction for each ARMS. ARMS retrieved in 2018 were replaced by uncolonized ARMS during the same dive. This new set of ARMS was retrieved and dismantled in 2019 (Table S1). In the present study, these samples were exclusively analyzed with metabarcoding under the new five structural categories scheme, with one DNA extraction and metabarcoding analysis for each category in each of the three ARMS per site.

### 2.2. Photo analyses

We used a SONY DRC-RX10M3 camera mounted on a fixed arm to photograph each face of every ARMS unit. Images were taken with a focal length of 35 mm, aperture F/6.3, exposure time 1/100 s, and ISO 80 sensitivity. Photographs were captured under optimal natural light conditions. Each image was 4864 × 3648 pixels (350 ppi, 24-bit color depth, sRGB color space). Images were subsequently cropped in IrfanView without quality loss and uploaded to the CORALNET website (https://coralnet.ucsd.edu/; Beijbom et al., 2015). Final cropped image files ranged from a few to about 10 MB, depending primarily on the proportion of uncolonized substrate.

Each image was then subdivided in 16 sub-squares with 4 randomly drawn points, to a total of 64 points (stratified random sampling, Figure 1b). For the annotation process, a list of labels was prepared (i.e. names of taxa or names of non-taxonomic categories to which a point could be assigned, fully described in Supplementary File S1). During the annotation process, this list has been modified to add new taxa and to refine categories. For labels added to distinguish two categories which were previously merged in a single label, experts had to update already assigned images. The CORALNET search engine helped finding the corresponding images.

Annotations were done manually until the automatic classifier was trained enough to propose labels, which were then either validated or corrected by experts (Beijbom et al., 2015; Chen et al., 2021; Williams et al., 2019). An initial model was generated after 50 annotated images, and a new model was trained every 50 additional annotated images. Over the course of 595 annotated images, 11 models were generated, with a maximum average accuracy of 69%. However, all label assignments were reviewed by experts, and the AI system served only as a tool to assist in the annotation process.

### 2.3. Metabarcoding protocol: from benchtop to desktop

Sessile material scraped from ARMS plate faces was homogenized (∼15 sec. in a blender), briefly rinsed, and transferred to clean 50 mL Falcon tubes filled with fresh 96 % ethanol. For DNA extraction, 1.5 mL of homogenate was used. To minimize cross-contamination, extractions were performed in individual tubes rather than in the 96-well plate format of the extraction kit (see Corse et al. 2017). DNA extractions and COI metabarcoding followed the protocol described by Thomasdotter et al. (2023), with the following key steps: (i) DNA was extracted using the NucleoSpin® Soil kit (Macherey-Nagel, Germany); (ii) the cytochrome c oxidase subunit I (COI) gene was amplified in triplicate PCRs with primers IIICRrevN (3[-GGNTGAACNGTNTAYCCNCC-5[) and HBR2d (5[-TAWACTTCDGGRTGNCCRAARAAYCA-3[), which include a heterogeneity spacer and 11–13 nucleotide sample-identifying tags, and target a broad spectrum of eukaryotic and algal phyla; (iii) a two-step tailed PCR generated paired-end, ready-to-load libraries; (iv) libraries were sequenced on an Illumina MiSeq using v2 chemistry (2 × 250 bp); and (v) a full series of negative and positive controls (two distinct mock communities) was incorporated.

Before the workflow was published by Thomasdotter et al. (2023), we conducted optimization trials (Supplementary File S2) to assess how the choice of DNA extraction kit and polymerase affected sequencing yield and taxonomic breadth. Six ARMS-derived samples (four sessile, two mobile) were each extracted at least twice using two DNA extraction kits, and all extracts were amplified in triplicate with two polymerases, generating 72 uniquely tagged amplicons for direct comparison. These trials showed that the NucleoSpin® Soil kit (Macherey-Nagel, Germany) recovered broader non-animal diversity without reducing metazoan richness, while the Qiagen Multiplex enzyme (Qiagen, Germany) produced higher read counts and more validated ASVs, leading us to adopt this extraction–polymerase combination for subsequent COI metabarcoding. This optimized workflow (described by Thomasdotter et al. 2023) was then applied to all study samples and forms the laboratory protocol implemented in this study.

For bioinformatic processing we used VTAM v0.2.0 (Validation and Taxonomic Assignment of Metabarcoding data; González et al., 2023), a pipeline specifically designed to leverage the presence of technical replicates and control samples (including negative controls and mock communities). The core principle of VTAM is to optimize parameter values across multiple filtering steps to maximize the removal of false-positive occurrences in control samples while retaining all expected variants (i.e., no false negatives in mock samples and the minimal possible number of false positives). Once optimized, these parameters are applied uniformly to all samples within the same sequencing library and run, under the rationale that parameter values validated on control samples are also appropriate for the corresponding experimental samples.

Briefly, paired-end reads were merged, quality filtered, demultiplexed, and trimmed to remove sequencing tags and primers. Default parameters were used, except that the minimum and maximum lengths of merged and trimmed reads were set to 295 and 331 bp, respectively. Amplicon Sequence Variants (ASVs) were initially filtered to remove low-frequency noise using low-stringency default thresholds, excluding occurrences with low absolute or relative read counts within samples or across variants. Such occurrences likely originate from sequencing or PCR errors, tag-jumps, or minor cross-sample contamination. Optimal parameter values for these filters were then inferred from the control samples, after which the filtering procedure was repeated using the optimized parameters. Final parameter settings are provided in Supplementary File S3.

To ensure reproducibility among technical replicates, only ASV occurrences detected in at least two of the three replicates per sample were retained, and replicates exhibiting composition profiles strongly divergent from the others were discarded. Subsequent filtering steps included the removal of potential chimeras and pseudogenes. Taxonomic assignment was performed in VTAM against the COInr database (version 2022-05-06; https://zenodo.org/records/6555985), after removing Insecta sequences to avoid spurious assignment of marine ASVs within Arthropoda (Meglécz, 2023a). Assignments were made by the algorithm of mkLTG (Meglécz, 2023b), using a stepwise identity threshold decreasing from 100 % to 85 %, ensuring high-resolution identifications (genus or species) when close reference sequences were available and lower-rank assignments when references were missing.

### 2.4. Statistical analyses of diversity

#### 2.4.1. Assemblage datasets

For the photographic analyses, CORALNET output tables give, for each photography, the number of points assigned to each label. For most comparisons with metabarcoding, non-biological labels such as not colonizable or bare substrate were removed, as was the label unknown, which generally represents undetermined organisms and could bias biodiversity estimates by conflating distinct taxa or by duplicating categories. For each remaining biological label, we calculated a standardized abundance by dividing the number of annotated points by the total number of labelled points on the image. From these data we constructed three nested datasets: (i) PhoLab0, comprising all biological labels (38 in total); (ii) PhoLab2, in which several labels were merged to yield 34 broader biological categories; and (iii) PhoPhy, in which labels were pooled into nine higher-level taxa corresponding to six animal phyla (Annelida, Bryozoa, Chordata, Cnidaria, Mollusca, Porifera), two photosynthetic groups (Ochrophyta and Rhodophyta), and a single label representing Foraminifera (Supplementary File S1).

For metabarcoding, the filtered dataset was first analyzed at the level of molecular operational taxonomic units (mOTUs)—clusters of amplicon sequence variants (ASVs) generated with Swarm v3 (Mahé et al., 2022) at d = 7—and then aggregated into higher-rank taxa (34 classes or 19 phyla or equivalent ranks) after removing mOTUs without any phylum-level assignment. (Supplementary File S1). Taxonomic richness and relative abundance were calculated for each ARMS plate face and for both methods, grouping photographic labels and mOTUs at the phylum level. Coverage-based rarefaction and extrapolation curves were computed from incidence data (presence–absence across samples) with the package iNEXT, version 3.0.1 (Chao et al., 2014; Hsieh et al., 2016) for both photography and metabarcoding datasets. These curves estimate how diversity accumulates with sample number and provide asymptotic estimates of total richness. Taxonomic richness was expressed as the Hill numbers with parameter q = 0 (D_0_), the exponential of Shannon-Wiener entropy (Hill number with q = 1, D_1_), and the inverse of the Simpson concentration index (Hill number with q = 2, D₂, i.e. the sum of squared frequencies of each taxon).

#### 2.4.2. α-diversity

For each of the four datasets, α-diversity (species richness) was quantified as the number of distinct photographic labels or mOTUs using PRIMER 7 (Clarke & Gorley, 2015) with the PERMANOVA+ add-on (Anderson et al., 2008). To verify that the choice of richness metric did not affect cross-method comparisons, we also calculated Shannon and Simpson indices and the related (not identical) Hill numbers (q = 1, and q = 2, respectively); because these metrics were strongly correlated with simple richness, they were not discussed further. Concordance between methods was evaluated by computing Pearson and Spearman correlation coefficients for paired photographic and metabarcoding α-diversity values from the same plate-face samples. Relative-abundance barplots were generated with the ggplot2 package, version 3.5.2 (Wickham et al., 2016) using all photographic labels and metabarcoding mOTUs aggregated to higher-rank taxa. Linear regressions confirmed that correlations were consistent across the different diversity indices (data not shown). Correlation matrices and visualizations were generated with the corrplot package, version 0.95 (Wei et al., 2017). Analyses were performed at two spatial scales: the sample level (166 individual plate faces) and the site level (10 sites represented by the combined abundances of the 17 faces of their analyzed ARMS).

#### 2.4.3. β-diversity, geography and environment

For each dataset, β-diversity was assessed from pairwise community dissimilarities calculated with the Bray-Curtis index (after a square root transformation of standardized abundances) and, for comparison, the Sørensen index, which relies solely on presence–absence data and is sometimes preferred for metabarcoding as read numbers may imperfectly reflect biomass or abundance (Deiner et al., 2017; Hajibabaei et al., 2019). Mantel tests were used to evaluate correlations in community dissimilarity between photographic and metabarcoding data (using the package ape, version 5.8.1; Paradis et al., 2019) at both sample level (all individual plate faces) and site level (45 non-redundant site pairs). To examine spatial and environmental structuring, least-cost sea distances (km) were computed among sites and correlated with Bray-Curtis dissimilarities using Mantel tests.

Environmental variables used in this study were derived from E.U. Copernicus Marine Service Information products. Physical time series (1987–2019) were obtained from the MEDSEA_MULTIYEAR_PHY_006_004 reanalysis (Escudier et al., 2020), and biogeochemical time series (1999–2018) from the MEDSEA_ANALYSIS_FORECAST_BIO_006_008 reanalysis (Teruzzi et al., 2021). For the 12 ARMS anchor points (latitude, longitude), 15 variables were extracted from 180 monthly CSV files, including temperature (thetao, bottomT), salinity (so), eastward and northward current velocity (uo, vo), sea surface height (zos), mixed layer thickness (mlost), nitrate (nit), phosphate (pho), phytoplankton carbon biomass (pcb), chlorophyll (chl), primary production (ppn), dissolved oxygen (dox), partial pressure of CO₂ (pco), and pH (ph). For each anchor point, basic statistics (mean, median, maximum, minimum, range, and first and third quartiles) were calculated for all variables using data averaged vertically between 1 and 50 m, on 4 × 4 km grids, from the full temporal coverage of the physical (1987–2019) and biogeochemical (1999–2018) models. All computations were performed in GRASS GIS version 8 (GRASS Development Team, 2025).

Euclidean distances based on scaled environmental variables (Latitude, Longitude, mean_thetao, range_thetao, mean_bottomT, range_bottomT, min_BottomT, max_bottomT, so, uo, vo, zos, mlost, nit, pho, pcb, chl, ppn, dox, pco, ph) were used in Mantel tests and partial Mantel tests (i.e. after sea least cost distance was accounted for) to disentangle geographic and environmental effects. Permutational multivariate analyses of variance (PERMANOVA; PRIMER 7 with PERMANOVA+) were performed on all photographic and metabarcoding datasets (across three taxonomic aggregation levels each) with models including combinations of factors such as region, site (nested within region or alone), face type (9T, BO, BC, TO, TC), defined by the combination of orientation (top vs. bottom), and compartmentalization (open vs. closed) (Table 2). For metabarcoding data only, we further compared the ARMS units analyzed face-by-face (one per site) against the two companion ARMS per site whose sessile fractions were pooled by means of barplots of relative abundance and richness by phylum and region. We also regressed site-level β-diversity with the logarithm of geographic distance and with Euclidian environmental distances using Mantel and partial Mantel tests as above. R analyses used the version 4.2.0 (R Core Development Team, 2023).

**Table 2:**
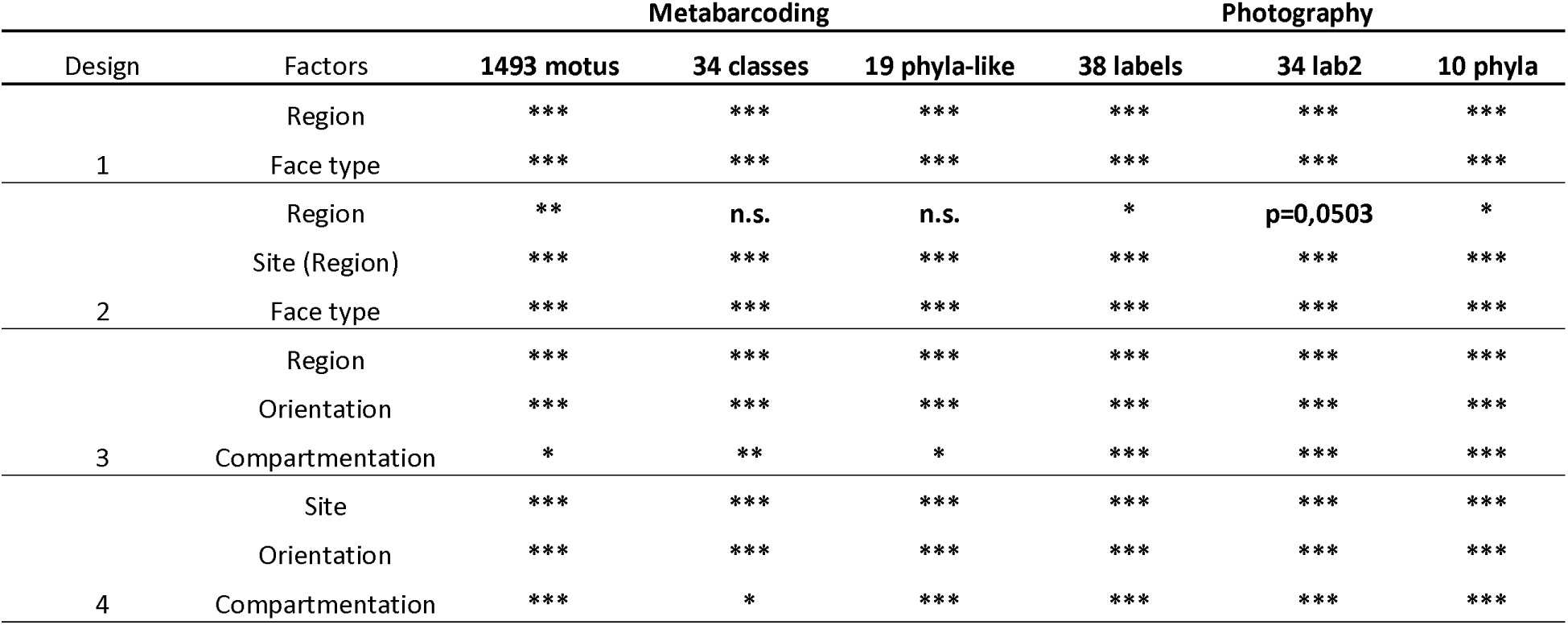
Main effects from PERMANOVA models. Summary of the primary factors explaining variation in community composition, as identified by permutational multivariate analysis of variance (PERMANOVA). The table reports the main effects retained in the final models for both photographic and metabarcoding datasets, with associated test statistics and significance levels using symbols (*** (p<0.001), **(p<0.01), *(p<0.05)), unless specified.

## 3. Results

### 3.1. Taxonomic composition, rarefaction patterns and face-type contrasts

Photography and metabarcoding revealed distinct yet partially overlapping characterization of ARMS sessile communities. To ensure robust cross-method comparisons, we removed four low-diversity samples—one photographic sample lacking any biological label (BAN3-3_5T) and two metabarcoding samples with very few reads and variants (BAN2-3_T9 and VIL2-1_B8). In the photographic dataset, non-biological categories dominated: “bare substrate” accounted for ∼43 % of the surface of all 598 images, “not colonizable” for ∼12 % (reflecting ARMS structural elements such as compartments and screws), and “detritus” for ∼5 %. Comparisons were therefore restricted to the 38 biological labels (algae, animals and Foraminifera) distributed across nine to ten phyla (Supplementary File S1).

Metabarcoding of the remaining 166 paired samples detected 1,493 mOTUs, an order of magnitude higher than the 38 photographic labels. Out of these ∼1,500 mOTUs, 150 were confidently assigned to 139 species, several of which were also observed visually (e.g. Botryllus schlosseri, Dictyota spp., Halopteris filicina, Sphaerococcus coronopifolius). Conversely, some solitary ascidians (e.g. Ciona spp.) were visible in photographs but absent in metabarcoding. As many marine organisms are missing from reference barcoding databases and since COI primers cannot amplify non-target templates, most mOTUs could not be confidently assigned below phylum level (Collins et al., 2019; Hintikka et al., 2022; Mugnai et al., 2021).

Sample richness ranged from 1 to 13 biological labels for photography and from 7 to 76 mOTUs for metabarcoding, underscoring the much finer taxonomic resolution of DNA data. Coverage-based rarefaction and extrapolation (Figure 2) confirmed this pattern: photographic richness approached saturation, whereas metabarcoding curves remained far from asymptotic, especially for q = 0 (rare taxa). Asymptotic Hill numbers of order 0 (richness), order 1 (abundant taxa, related to Shannon’s entropy) and order 2 (dominant taxa, related to Simpson’s index) were 40 (s.e. = 6.9), 20.4 (s.e. = 0.5), and 16 (s.e. = 0.44) labels for photography, compared with 2 391 (s.e. = 75), 902 (s.e. = 16), and 455 (s.e. = 10) mOTUs for metabarcoding.

**Figure 2:**
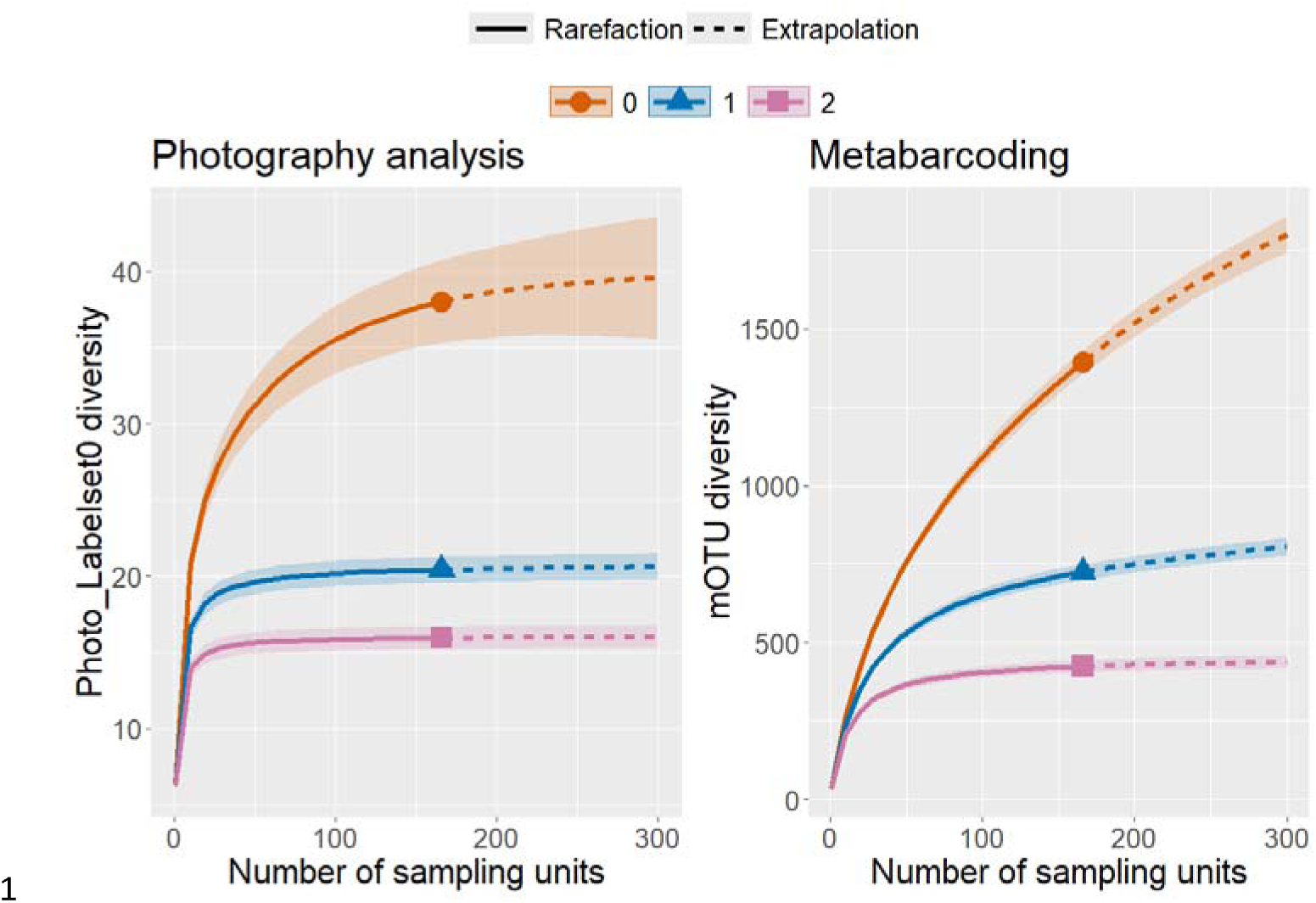
Taxa rarefaction and extrapolation curves. The number of photographic taxonomic labels or metabarcoding molecular operational taxonomic units (mOTUs) is plotted on the vertical axis as a function of the number of photographic or metabarcoding samples on the horizontal axis. Curves represent three types of Hill numbers defined by the q parameter: taxonomic richness (q = 0), i.e. the number of labels or mOTUs (orange); Shannon-like diversity (q = 1), i.e. the exponential of Shannon entropy (blue); and Simpson-like diversity (q = 2), i.e. the inverse of Simpson concentration (pink). Shaded areas denote standard-error confidence intervals. The metabarcoding dataset here corresponds to the 17-faces design (for comparisons with photography).

Taxonomic distributions also differed between methods and among plate faces (Figure 3). Both approaches revealed higher Rhodophyta abundance on top faces, but Bryozoa and Annelida were proportionally far richer in photographs, whereas Arthropoda and Echinodermata were detected only by metabarcoding.

**Figure 3:**
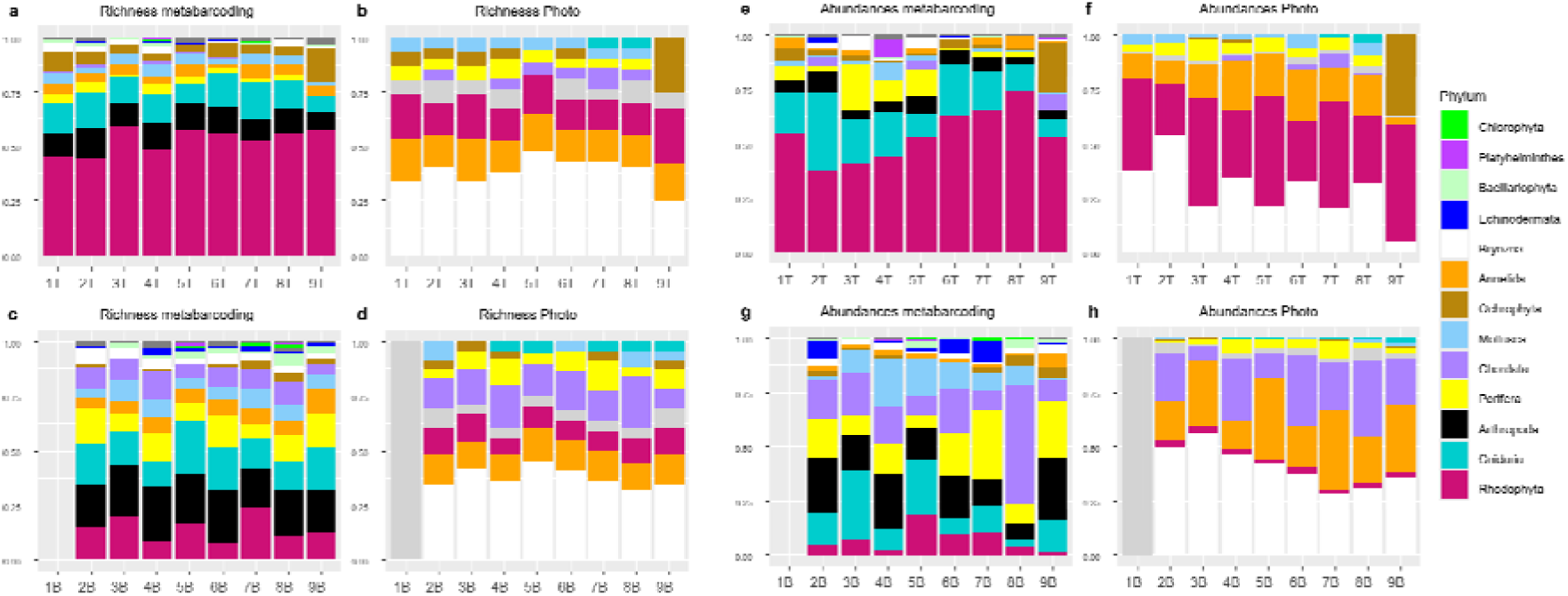
Barplots of taxonomic composition by phylum for each method and plate face. Top and bottom plate faces are shown in the upper (a, b, e, f) and lower (c, d, g, h) panels, respectively. Relative richness (a, b, c, d) and relative abundance (e, f, g, h) are presented for metabarcoding (a, c, e, g) and photography-based analysis (b, d, f, h). Note that Chordata consist mainly of Ascidiacea rather than fishes, which were absent from the photographic data and relatively rare in the metabarcoding results. Unassigned mOTUs are not displayed, and all mOTUs belonging to sparsely represented phyla or higher-rank taxa are pooled in dark grey, a category not shown separately in the legend. The taxa in the unique legend are ordered by their abundances in metabarcoding. The metabarcoding dataset used here corresponds to the 17-faces design (for comparisons with photography).

Pooling all sessile fractions per ARMS drastically reduced diversity in metabarcoding: 22 pooled samples (two ARMS per site plus one biological replicate for MAR3 and MAR4) yielded 559 mOTUs—less than half of the 1,493 mOTUs obtained from 166 separated-face samples from ten ARMS (Figure 4). Phylum abundances were more even in the face-by-face design, whereas pooled samples over-represented Ochrophyta and, to a lesser extent, Porifera.

**Figure 4:**
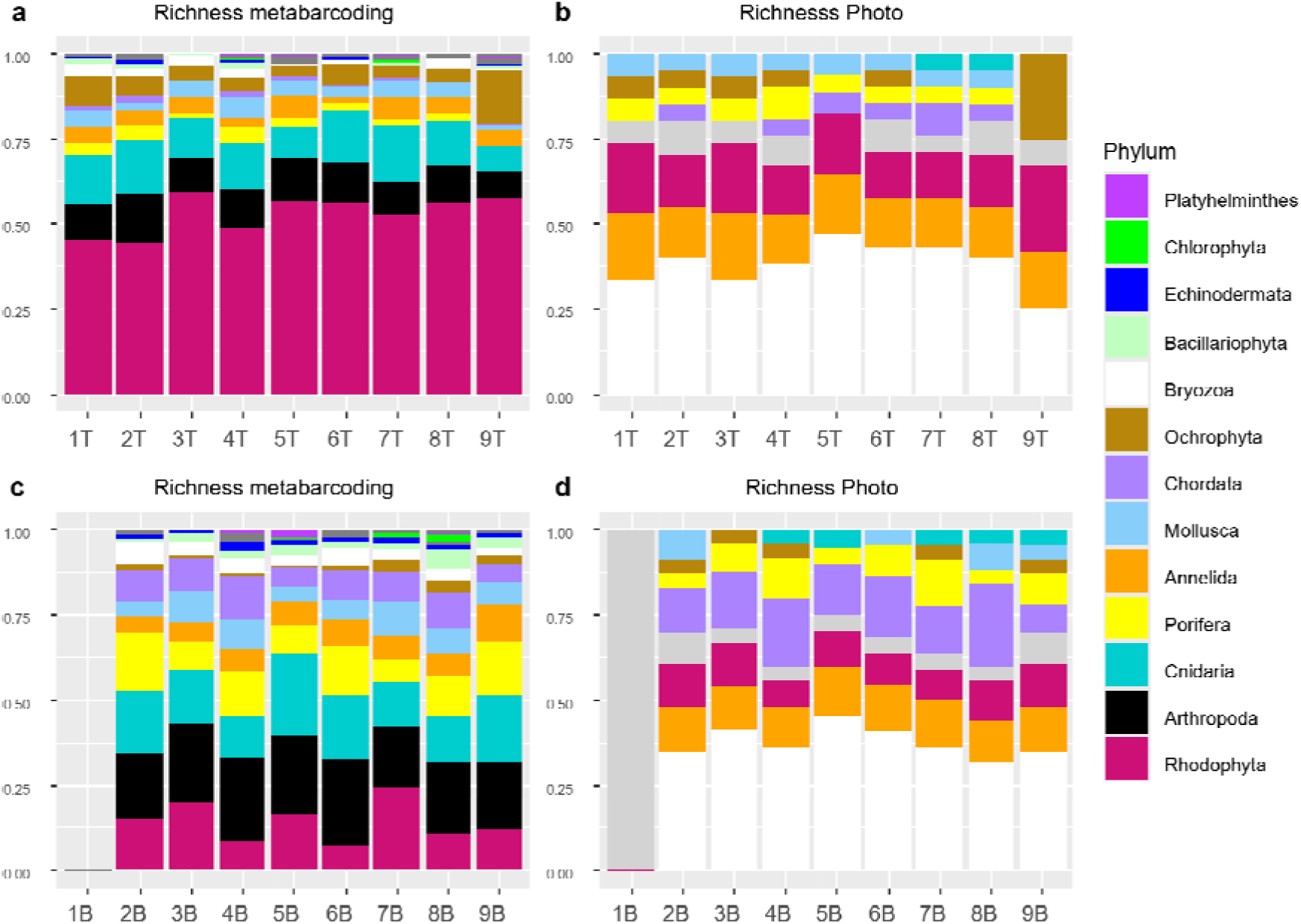
Comparison of metabarcoding designs: separate versus pooled ARMS faces. Bar plots show, for each site, the relative abundance of mOTUs by phylum. Top panel: each bar represents the combined total of 17 individual face samples from a single ARMS (with minor exceptions where some faces were discarded; see text). Bottom panel: each bar represents the combined total of two pooled sessile-fraction samples, each derived from a whole ARMS, following the standard NOAA protocol. This dataset corresponds to the ARMS retrieved in 2018 only.

In the 2019 metabarcoding dataset, based on five structural face categories accross three ARMS per site, 137 samples were retained after quality filtering (13 removed from 150). These yielded 1,541 distinct mOTUs, slightly exceeding the 1,493 recovered from the 166 individual-face samples analyzed in 2018.

### 3.2. Relationships between photographic and metabarcoding diversity

Sample-level α-diversity values showed a slight but significant negative correlation between photographic and metabarcoding datasets (Pearson R = –0.17, p = 0.0245; Figure 5). After pooling all plate-face samples within each site, the relationship became positive but non-significant (R = 0.24, p = 0.51), insignificance being likely driven by an outlier site from the Villefranche-sur-Mer region. Site richness was roughly 15-fold higher for metabarcoding than for photography (Figure 5). When taxa were aggregated to higher ranks, positive site-level correlations became clearer. The strongest correspondence (Figure S1) occurred when metabarcoding data collapsed to 34 classes were compared either with the full photographic label set (38 labels; R = 0.71, p = 0.021), or with the intermediate labelset2 (34 labels; R = 0.69, p = 0.026; Figure S1). In contrast, β-diversity (Bray-Curtis dissimilarity) exhibited consistently positive and stronger correlations between methods, both at the sample level (R = 0.39, Mantel p = 0.0001; Figure 5) and at the site level (R = 0.72, Mantel p < 0.0001; Supplementary Figure S1). These positive correlations persisted at all taxonomic aggregation levels, but coefficients declined steadily with coarser taxonomic ranks, reaching non-significant values (R ≈ 0.20) when both datasets were reduced to phylum level (Supplementary Figure S1). The contrasting signs of α-diversity correlations at sample versus site level largely can be attributed to strong differences among ARMS face types: for instance, the superior 9T face was the most species-rich in metabarcoding but among the least diverse in photographic data, whereas bottom-open (BO) faces showed the opposite trend (Figure 6).

**Figure 5:**
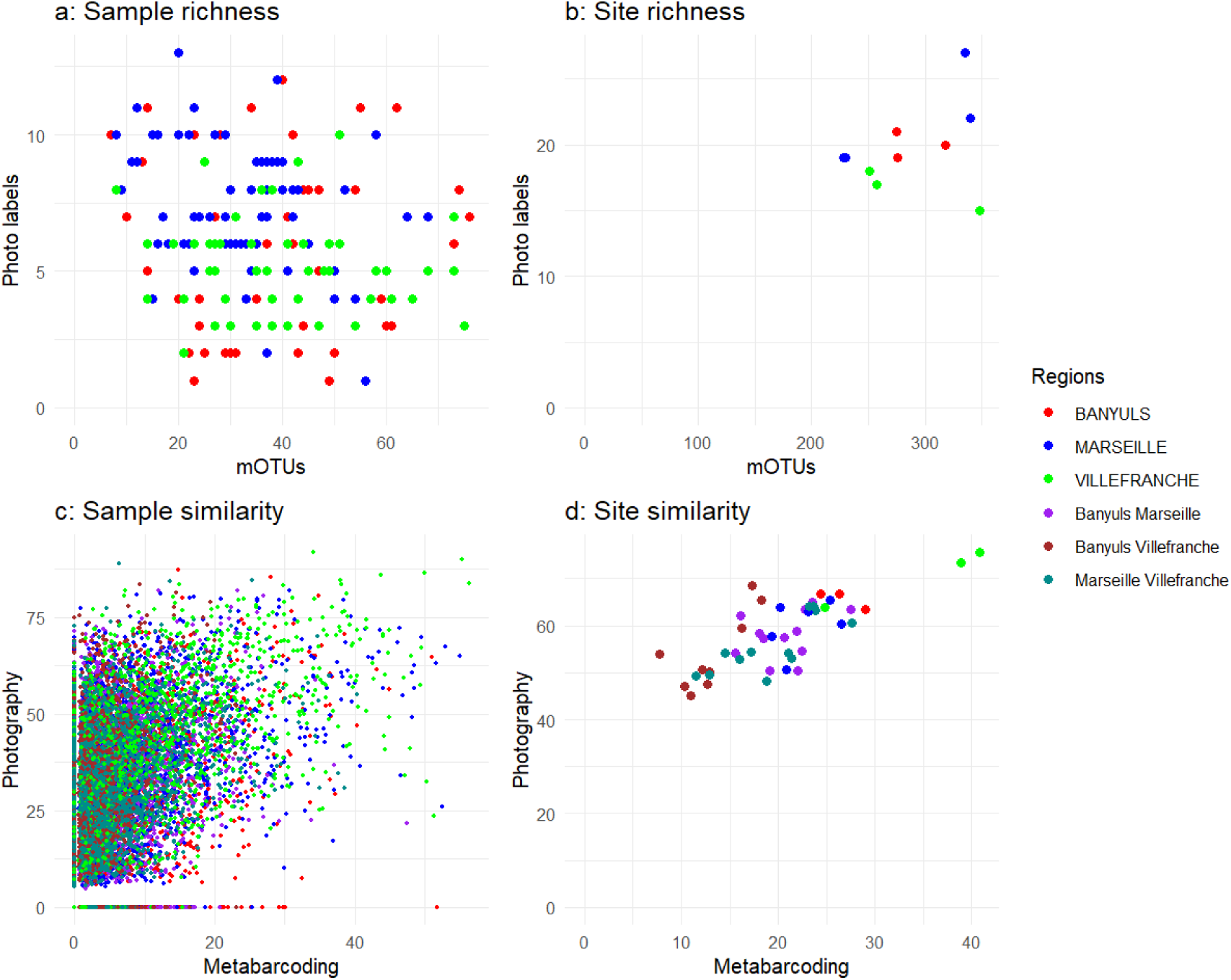
Correlation of diversity measures between photographic and metabarcoding methods (lowest taxonomic ranks). Horizontal axes show values derived from metabarcoding data (1,393 mOTUs), and vertical axes show values derived from photographic data (38 labels, labelset 0). Upper panels (a, b) present α-diversity (richness), and lower panels (c, d) present β-diversity (Bray–Curtis similarity) for pairwise comparisons. Left panels (a, c) display sample-level data, whereas right panels (b, d) display site-level data. The metabarcoding dataset used here corresponds to the 17-faces design (for comparisons with photography).

**Figure 6:**
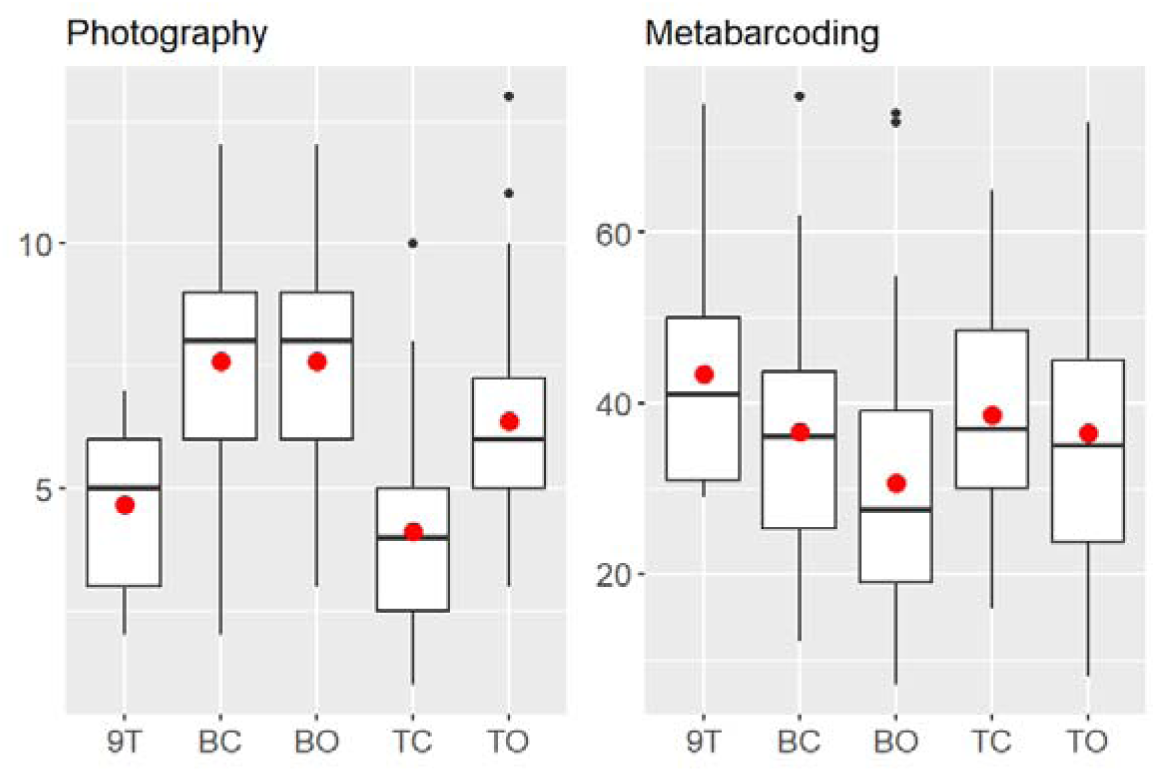
Taxonomic richness across five ARMS structural face categories and both survey methods. Left panel: photography-based analysis (vertical axis: number of distinct biological labels). Right panel: metabarcoding (vertical axis: number of molecular operational taxonomic units, mOTUs). Red dots indicate the mean; thick horizontal bars show the median; box limits correspond to the first (Q1) and third (Q3) quartiles; whiskers extend to the most extreme data points within 1.5 × IQR (interquartile range); and isolated points represent outliers beyond this range. Structural face categories that are least diverse in photographic analyses (9T, TC) tend to be the most diverse in COI metabarcoding. The metabarcoding dataset used here corresponds to the 17-faces design (for comparisons with photography).

### 3.3. Geographical and ecological effects on β-diversity

Both photographic and metabarcoding datasets revealed strong spatial and ecological structuring of community composition. PERMANOVA models (Table 2) detected highly significant effects of region, site, and plate-face type on β-diversity. When region and site (nested within region) were included jointly, regional effects weakened (higher p-values), indicating that much of the regional signal was captured at the site level. Face orientation (top vs. bottom) was consistently highly significant, whereas (open vs. closed) was strongly significant for photographic data and moderately significant for metabarcoding. Non-metric multidimensional scaling (nMDS) ordinations (Figure 7) corroborated these patterns: samples separated clearly by face orientation, open–closed differences were more diffuse, and regional structure appeared as an east–west gradient along a diagonal axis. This geographical gradient was slightly more pronounced for metabarcoding, whereas fine-scale ecological differentiation within-ARMS (related to light and microhabitat) was clearer in the photographic ordination. Analyses based on the Sørensen index, which is based on presence–absence rather than abundance, yielded very similar results (data not shown).

**Figure 7:**
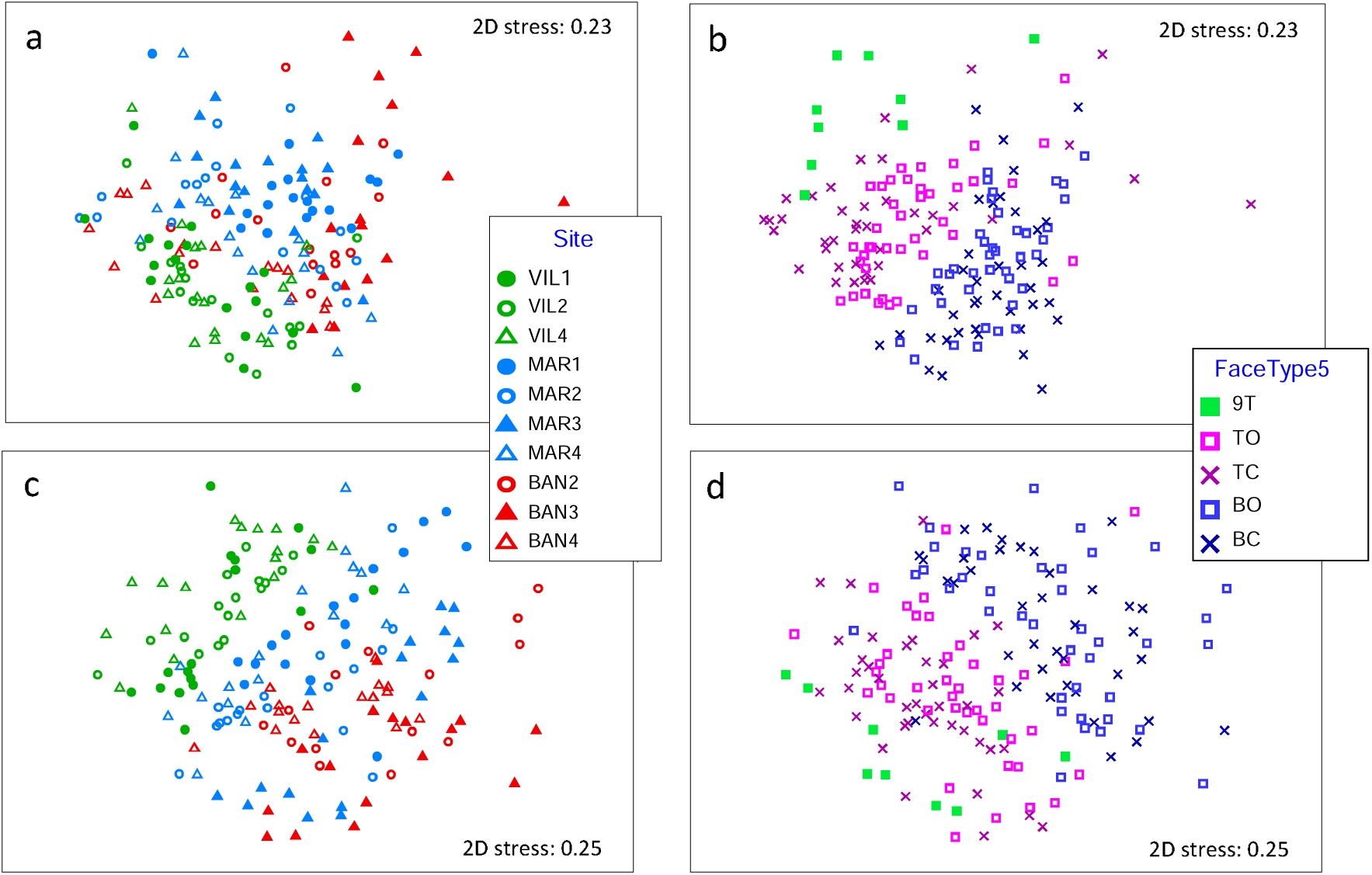
β-diversity comparison using non-metric multidimensional scaling (NMDS). Ordination of samples based on Bray–Curtis dissimilarity computed from standardized and square-root–transformed abundances (read numbers for metabarcoding; surface cover for photography). Top panels (a, b): photography data (38 labels, labelset 0). Bottom panels (c, d): metabarcoding data (1,493 mOTUs). Left subpanels (a, c): points coloured by site and by region (green = Villefranche, blue = Marseille, red–pink = Banyuls). Right subpanels (b; d): points coloured by face type, with symbols indicating compartmentation (× = closed faces; open squares = open faces) and orientation (blue = bottom; pink = top; light green = 9T faces receiving more surface light). The metabarcoding dataset used here corresponds to the 17-faces design (for comparisons with photography).

Community similarity declined significantly with the logarithm of sea least-cost geographic distance for both methods, but more tightly for metabarcoding (Mantel R = 0.70, p = 0.00003) than for photography (R = 0.55, p = 0.00038; Figure 8 a,b). The weaker relationship for photography likely reflects unusually high similarity between the most distant regions (Banyuls-sur-Mer and Villefranche-sur-Mer) and low similarity among some Marseille sites. Taxonomic aggregation reduced distance–decay correlations for both datasets, though correlations remained significant. When metabarcoding mOTUs were collapsed to class or phylum, Mantel coefficients decreased to R = 0.43 (p = 0.00183) and R = 0.34 (p = 0.00482), respectively (Supplementary File S5). Similarly, aggregating photographic data to labelset2 or phylum yielded R = 0.47 (p = 0.00138) and R = 0.27 (p = 0.01593) (Supplementary File S5). By contrast, pooled-faces metabarcoding performed worst, producing the lowest and non-significant correlations with geographic distance (R = 0.18, p = 0.1359; Figure 8c).

**Figure 8:**
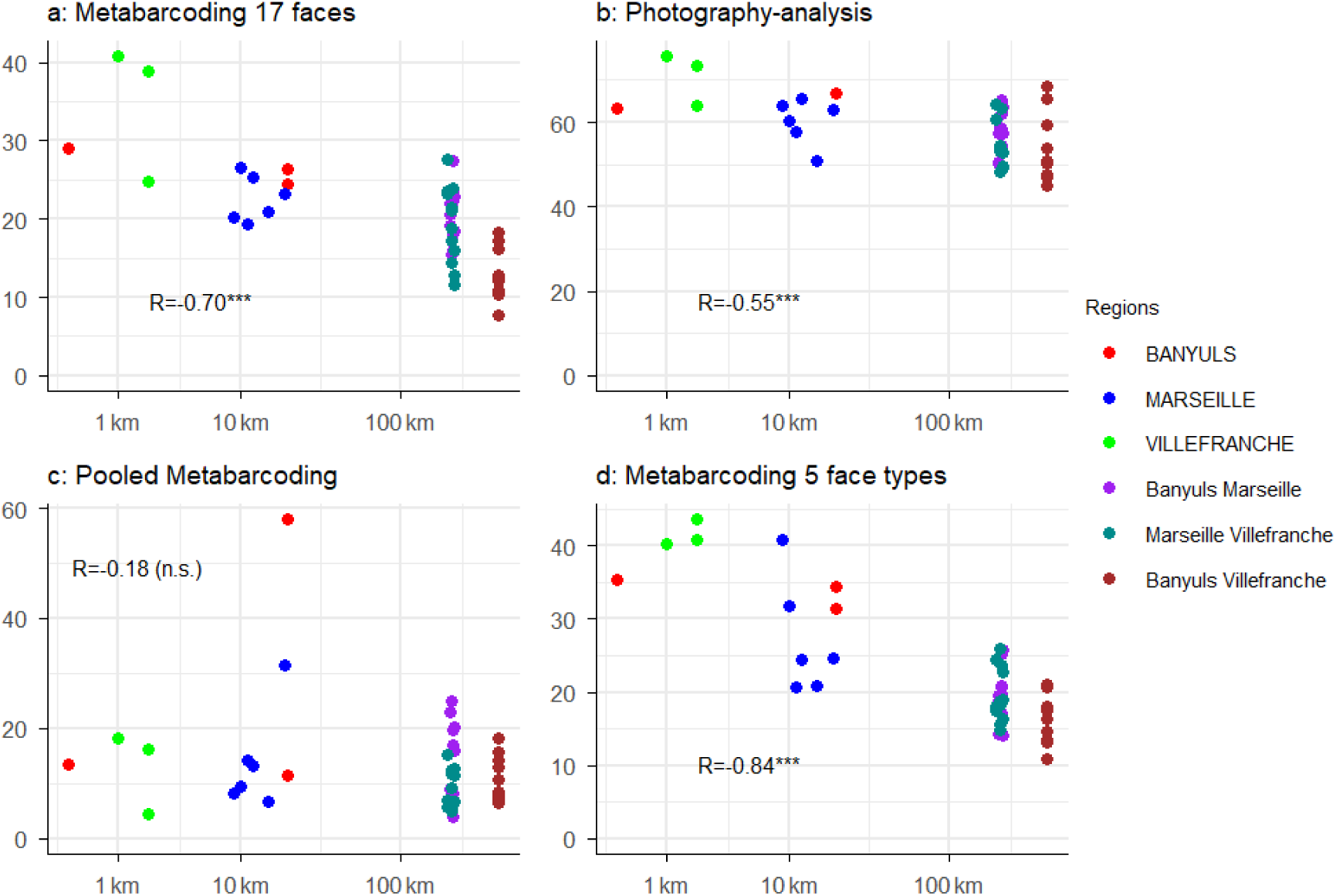
Community similarity versus geographic distance. Scatterplots of Bray-Curtis dissimilarity (vertical axis) against sea the least-cost geographic distance (km) (horizontal axis, logarithmic scale). Metabarcoding data—(a) separate faces (166 samples before pooling by site, retrieved in 2018), (c) pooled sessile fractions (22 samples before pooling by site, retrieved in 2018), and (d) five sessile fractions (137 samples before pooling by site, ARMS retrieved in 2019). Right panel (b): photography-based data (2018). Pearson correlation coefficients and significance levels are indicated on each plot. Note: the vertical axis scale varies among panels.

Metabarcoding based on five structural face categories (three ARMS per site, 2019 dataset) yielded the strongest distance–decay relationship (R=0.84, p=0.00002; Figure 8d), outperforming the 17-face design despite using fewer samples (137 vs. 166). Regression intercepts indicated that similarity at 0 km averaged 64 % for photography, 27 % for face-by-face metabarcoding, and only 16 % for pooled metabarcoding, highlighting the reduced consistency of pooled samples. The greater distance of metabarcoding rarefaction curves from their asymptotes likely contributes to this pattern, indicating higher sampling variance relative to photography.

Environmental variables also helped explain β-diversity patterns (Supplementary File S4). Mantel tests showed that scaled Euclidean distances in several environmental descriptors were significantly correlated with community dissimilarity, although geographic distance consistently explained more variance. Partial Mantel tests controlling for geography identified four significant predictors for metabarcoding (eastward current velocity, nitrate, phosphate, and primary production) and two for photography (nitrate and chlorophyll concentration). In addition, chlorophyll was nearly significant for metabarcoding, and eastward current velocity, phosphate, phytoplankton carbon biomass, and primary production approached significance for photography. Metabarcoding using five structural face categories (three ARMS per site) identified six significant environmental variables, the highest number across all datasets.

## 4. Discussion

Our key methodological advance—separating ARMS faces for metabarcoding—proved decisive for improving biodiversity assessments. By increasing detected richness, reducing sampling variance, and strengthening biogeographic and environmental correlations, this approach markedly outperformed the standard NOAA pooling protocol and provides a new benchmark for standardized eDNA monitoring. In the following discussion, we examine (i) the complementarity between photography and metabarcoding, (ii) the causes of their differing performance, and (iii) the implications of our results for sampling design and for routine monitoring frameworks—with a focus on the benefits of separating ARMS structural face categories (or all 17 faces) compared with pooled designs for metabarcoding. Finally, we consider the broader implications of these findings for long-term marine biodiversity monitoring and management.

### 4.1. Complementarity of photography and metabarcoding

α- and γ-diversity were markedly higher in metabarcoding than in photographic analyses, even under a stringent bioinformatic pipeline (González et al., 2023). This pattern reflects the finer taxonomic resolution of COI metabarcoding—which can distinguish cryptic species lacking morphological differentiation (Cahill et al., 2024; Chenuil et al., 2019)—as well as its ability to detect a broader size range of organisms. The two methods also differed in how they resolved variations among ARMS structural face categories. Photography recorded the lowest diversity on superior faces (9T), whereas these yielded the highest richness by metabarcoding. A likely explanation is that dense macroalgal cover (e.g. Ochrophyta such as Dictyota spp.) obscures underlying microhabitats from visual detection, while providing more colonizable surface for cryptic taxa that are well captured by DNA. Conversely, photography revealed the lowest richness on top-closed faces (a pattern not observed in metabarcoding) likely reflecting differences in light exposure and the availability of bare-substrate. These contrasting face-type effects explain the negative correlation of α-diversity between methods at the sample level, whereas pooled site-level richness showed a positive—but non-significant—correlation, possibly constrained by the limited number of sites (n = 10)

The taxonomic spectra recovered by the two methods overlapped but differed in detail, highlighting their complementarity. Every phylum recorded by photography was also detected by metabarcoding, yet groups such as Bryozoa and Annelida were abundant in photographs but rare in metabarcoding. Conversely, most of the metabarcoding diversity consisted of mOTUs without phylum-level assignments, either because reference sequences are lacking for these lineages or because the amplified COI fragments derive from poorly catalogued or non-metazoan organisms, including unicellular eukaryotes and even prokaryotes (Collins et al., 2019; Mugnai et al., 2021; Hintikka et al., 2022). Although photographic analysis detected far fewer taxa than metabarcoding, both approaches effectively revealed significant effects of all tested spatial and ecological factors on community composition. These factors included region (three regions separated by up to ∼395 km), site within region (0–20 km apart), and ARMS face type, defined by the combination of orientation (top vs. bottom) and compartmentation (open vs. closed), resulting in five categories (9T, top open, top closed, bottom open, bottom closed). However, subtle but consistent differences emerged in the drivers of community composition detected by each method, p-values differed slightly between them, indicating nuanced contrasts in sensitivity. Photography detected more significantly than metabarcoding difference in microhabitat within ARMS (plate compartmentation).

Together, these comparisons show that photography and metabarcoding provide complementary insights into marine hard-bottom biodiversity. Photography offers direct visual estimates of macroscopic cover, especially macroalgae, while metabarcoding captures cryptic, microscopic, or early-stage organisms and preserves fine-scale heterogeneity at the scale of individual plate faces. In future monitoring frameworks, the balance between both approaches may be tailored to specific management goals and ecosystem types: photographic analyses may best track dominant, visually identifiable taxa or estimating surface cover, whereas metabarcoding may be preferable for detecting cryptic assemblages, early colonizers, or biodiversity shifts in turbid, structurally complex, or species-rich environments. The broad convergence between methods supports their reliability and robustness, and the stronger geographic and environmental signal in metabarcoding highlights its greater sensitivity to subtle compositional change.

### 4.2. Taxonomic resolution explains the high sensitivity of metabarcoding to environmental and spatial gradients

In contrast with α-diversity patterns at sample levels, β-diversity patterns were strongly congruent between methods, with significant Mantel correlations at both sample and site levels (Figure 5, Figure S1). This indicates that both approaches reliably captured community structure. However, metabarcoding revealed stronger correlation of community similarity with geographic distance (R = 0.70 vs. 0.55 for photography, with a single ARMS per site). Likewise, both methods revealed similar environmental variables correlated with community dissimilarity, but metabarcoding consistently produced higher statistical power, with several variables that were marginally non-significant in photographic data becoming significant in metabarcoding. Aggregating mOTUs into higher ranks (i.e., classes and phyla) progressively weakened these correlations, ultimately falling below those from photography, demonstrating that the stronger biogeographic and environmental signal stems from its finer taxonomic resolution.

We expected mOTU-level metabarcoding to reflect geographic distances more clearly than photography. This is because metabarcoding can detect intraspecific diversity (e.g., Thomasdotter et al., 2023; Turon et al., 2020), and population genetic structure often reflects isolation-by-distance or oceanographic connectivity (Legrand et al., 2022). Marine species frequently exhibit spatial variation in mitochondrial COI haplotype frequencies, including along the French Mediterranean coast where the present study takes place (Boissin et al., 2015; Cahill et al., 2017; Penant et al., 2013). Moreover, numerous cryptic species complexes, genetically distinct but morphologically indistinguishable, exhibit clear COI divergence (Cahill et al., 2024; Chenuil et al., 2019; Egea et al., 2016). Photographic analyses cannot resolve such variation, and most visually distinguishable taxa using our photographic protocol are widespread throughout the north-western Mediterranean and therefore less likely to reflect geographic distances. Consequently, metabarcoding was expected to reflect geographic distances more clearly than photographic analyses.

For environmental factors, expectations differ. Because metabarcoding detects small, fast-responding organisms, including unicellular taxa amplified by COI, community changes might be captured more rapidly by DNA than by imagery. This aligns with the classical view that microorganisms respond more strongly to environmental conditions than to geographic barriers, a concept captured by the phrase “everything is everywhere, but the environment selects” (De Wit and Bouvier, 2006; Van der Gast, 2013). Under this paradigm, metabarcoding could be especially sensitive to environmental variation. However, when environmental gradients are spatially structured, the expected relative performance of imaging vs. metabarcoding becomes less obvious. Abundant taxa well adapted to local conditions should, in principle, have population densities that mirror environmental parameters more faithfully than simple presence–absence data. This would give photography an advantage because percentage cover is a meaningful ecological proxy. By contrast, metabarcoding read counts are imperfect indicators of abundance, owing to multiple sources of technical biases: (i) subsampling of the homogenized community may under-represent rare taxa; (ii) DNA extraction efficiencies varies among taxa, and (iii) PCR amplification is taxon-specific, with some lineages weakly or not amplified (Lamb et al., 2019; van der Loos and Nijland, 2021). All else being equal, such biases should reduce the sensitivity of metabarcoding to environmental gradients. Yet our results contradict this expectation: metabarcoding exhibited stronger correlations with environmental variables than photography. This indicates that the far greater taxonomic richness recovered by metabarcoding compensates for its lower quantitative fidelity, enhancing its ability to detect environmentally driven changes in community composition (see also: Deiner et al. 2017).

### 4.3. Implications and perspectives for routine monitoring

Standardized ARMS protocols are essential for generating DNA- and photography-based biodiversity data that remain comparable across studies and over time (Ransome et al., 2017). Harmonized procedures—such as those promoted by the NOAA protocol and now adopted by international initiatives like ARMS-MBON—have become a cornerstone of global marine biodiversity observation (Obst et al., 2020; Daraghmeh et al., 2025). Standardization is particularly critical for eDNA metabarcoding, because even minor changes in sampling, storage, or sequencing can cascade through bioinformatic workflows and compromise inter-study comparability (Zinger et al., 2019; van der Loos & Nijland, 2021; Alberdi et al., 2018). Building on this recognized need, our study refines key steps of ARMS-based monitoring—from field deployment to molecular and photographic processing—so as to strengthen reproducibility and long-term comparability, providing the baseline on which more detailed sampling strategies can build.

#### 4.3.1. Sampling design and sampling variance: why pooling all ARMS faces before metabarcoding should be avoided

A major difference between the four methods implemented in this study concerns sampling variance. Metabarcoding is more prone to stochasticity than photography, consistently missing a larger fraction of taxa that are in principle detectable. In our dataset, this was evident from the markedly lower community similarity between adjacent sites and from the rarefaction curves further from their asymptotes compared with photographic data. Such elevated among-sample variability accords with earlier reports of eDNA detection heterogeneity (Zinger et al., 2019; Alberdi et al., 2018; Lamb et al., 2019). The elevated variance observed in metabarcoding is not merely a statistical consequence of detecting more numerous taxa, which would naturally increase sample-to-sample variability if sample size were limited. Sequencing depth was not limiting in our study (not shown), suggesting that the observed variance reflects technical artefacts introduced at multiple stages of the workflow, including DNA subsampling, extraction efficiency, PCR bias, and data filtering (Lamb et al., 2019; van der Loos & Nijland, 2021; Zinger et al., 2019). By contrast, photographic analysis—although based on 64 randomly drawn points per image and thus theoretically subject to some sampling errors—missed far fewer taxa and yielded more consistent estimates of local similarity. These results strongly argue against pooling all sessile fractions from a single ARMS prior to metabarcoding. Indeed, the similarity of adjacent sites reached values as low as 5% in pooled faces metabarcoding, which is extremely low as compared to a minimum of 20 % for separated-faces metabarcoding, and 50 % for photographic analyses.

Analyzing the 17 faces of a single ARMS per site separately increased the number of detected mOTUs by more than 2.4-fold compared with pooling all sessile fractions from 22 ARMS (across ten sites). The gain in diversity was even higher with the design separating five structural face categories across three ARMS per site. In addition to reducing sampling variance, separating structural face categories (e.g. top vs bottom, open vs closed) likely also limits the loss of diversity caused by DNA degradation or by PCR inhibition from chemical compounds produced by certain sessile taxa—an adaptation for space competition typical of sessile organisms—that are unevenly distributed across structural face categories, as reported for sponges (Leray & Knowlton, 2015). This interpretation is further supported by our observation of shifts in phylum representation between pooled and face-by-face designs, suggesting that DNA from distinct phyla is not extracted and/or amplified with equal efficiency depending on the taxonomic composition of the mixture (see also: Lamb et al., 2019; van der Loos & Nijland, 2021). This hypothesis—namely that separating samples into structural face categories improves site characterization beyond the mere reduction of sampling variance—is also supported by the observation that community composition based on the five structural face categories captured geographic and environmental variation more strongly than the 17-face design, even though it was based on fewer samples (137 vs. 166 across ten sites). We thus demonstrate that, beyond increasing detected diversity, sampling design has a decisive impact on the ability to identify ecological drivers of community composition, which is critical for effective monitoring.

Our findings challenge the standard NOAA protocol, which recommends three ARMS units per site with pooled sessile fractions. Given the very low correlations we observed between community differentiation and geographic or environmental distances in pooled designs (Figure 8), it is unlikely that NOAA protocol (based on three pooled ARMS per site) would perform substantially better than the two pooled ARMS per sites analyzed here. Because separating 17 sessile samples per ARMS entails substantial extra work, and because the five-category scheme provided an even clearer ecological signal, we subsequently adopted the intermediate solution for the SEAMoBB project: pooling faces into five structural categories based on orientation and compartmentalization, while retaining three ARMS per site. We considered including plate number (i.e., distance from the seafloor) as an additional structural factor. However, preliminary analyses showed no consistent pattern (data not shown), and plate number cannot be fully crossed—with respect to statistical design—with orientation and compartmentalization. Overall, this modified protocol offers a balanced compromise between logistical effort and ecological resolution.

#### 4.3.2. Refining photographic analysis: greater precision with minimal additional effort

Photography-based analyses also offer room for methodological refinement. Most ARMS image studies have relied on 50 random points per plate face, following a comparative analysis indicating that this density is generally sufficient. By increasing the density to 64 points, we improved precision without a meaningful increase in processing time, and found that photographic data were much less prone to sampling variance than metabarcoding, because adjacent sites harbored more similar communities (see also David et al., 2019). Although we did not quantify the gain with respect to sampling 50 points, we experienced that the working time was comparable when adding 14 points, while abundances are more reliably estimated. Nevertheless, further gains are likely to come from improving image quality and resolution, which would require higher-grade cameras and greater digital-storage capacity, rather than from further increasing point density. Expanding the number of taxonomic labels can enhance detection of changes in community structure, biomass and anthropogenic impacts (PERMANOVA tests, not shown), but it also increases training time and inter-expert variability (see also Beijbom et al., 2015; Williams et al., 2019). Looking ahead, artificial-intelligence–assisted image recognition, coupled with steadily increasing computing power, is expected to accelerate point annotation, improve consistency, and reduce expert bias (Bravo et al. 2021; Raphael et al., 2020; Trotter et al., 2025), thereby further enhancing the efficiency and reproducibility of ARMS imagery.

#### 4.3.3. Traceable, quality-controlled bioinformatics

A key requirement for long-term eDNA monitoring is to generate datasets comparable across sites, seasons, and sequencing runs—a transition from research to operational biomonitoring that requires rigorous quality control and standardization (Pawlowski et al., 2018; Petit-Marty et al., 2023; Rimet et al., 2025). International standardization bodies (ISO/CEN; Pawlowski et al., 2018) and recent reviews (Leese et al., 2018; Hakimzadeh et al., 2024; Rimet et al., 2025) specifically call for bioinformatic protocols that minimize false positives and false negatives, ensure full traceability from field sampling to biodiversity indices, and enable reproducible, large-scale inter-comparisons. The bioinformatic considerations raised by Turon et al. (2025) reinforce these requirements, showing that temporal trends can be confounded by technical discrepancies across sequencing runs or workflows and thus emphasizing the need for control-driven, fully traceable pipelines.

In this context, the VTAM pipeline proved crucial as it explicitly integrates negative controls, mock communities, and technical replicates to optimize filtering parameters, thereby reducing false positives and false negatives and ensuring full traceability at every filtering step (González et al., 2023). By adopting this validated pipeline, we ensured that the strong biogeographic and environmental patterns revealed by our face-by-face metabarcoding reflect true biological signals rather than analytical artefacts. This conclusion is further supported by recent studies demonstrating that VTAM-filtered metabarcoding can generate highly validated data for fine-scale, hypothesis-driven analyses of community assembly and ecological interactions across aquatic ecosystems (e.g. Esposito et al. 2022; Ruiz et al., 2025; Villsen et al., 2025). VTAM thus provides a robust and transparent framework for standardizing ARMS metabarcoding data, facilitating inter-study comparisons and ensuring that biodiversity signals reflect genuine ecological patterns. This traceable and reproducible workflow directly meets the priorities identified in recent international assessments and underpins the interoperability required for future ARMS-based observatories.

#### 4.3.4. Costs and scalability

Cost and time considerations reinforce these conclusions. Assigning 64 points per image required only about 4–8 minutes for a trained analyst, so processing our 163 images amounted to less than one week of full-time work after an initial ten-day training period. By contrast, metabarcoding required roughly one month of benchtop labor for ∼160 samples and about €6 000 for reagents consumables and sequencing, half of which being for high-throughput sequencing. The recent adoption of Illumina NovaSeq sequencing has reduced per-sample sequencing costs more than tenfold, but exploiting these economies of scale requires pooling sufficient libraries to fill sequencing runs, which may introduce delays that must be balanced against monitoring frequency. Similar trade-offs between detection sensitivity and economic efficiency have been demonstrated in other eDNA studies, where laboratory costs can offset field savings and strongly influence optimal survey design (Smart et al., 2016). Broader reviews likewise stress that the choice of sampling and processing methods remains a major determinant of cost, throughput, and inter-study comparability in benthic metabarcoding (Pawlowski et al., 2022), underlining the need to balance detection sensitivity with cost efficiency when designing long-term observatories. Together, these cost and scalability insights can help define practical designs for long-term eDNA observatories, offering practical guidance for monitoring programs. While consumable and sequencing costs are steadily decreasing thanks to technological advances, the main long-term constraints for large-scale monitoring are more likely to stem from field efforts and human processing time. Optimizing this balance is therefore more critical than temporary reagent or sequencing prices, and sustainable implementation will depend on coordinated infrastructures and shared facilities to ensure cost-efficiency and dataset comparability across regions and monitoring programs.

## 5. Conclusion

By coupling high-resolution photographic validation with a traceable, control-driven metabarcoding workflow, this study refines ARMS-based eDNA monitoring into a reproducible and scalable framework for long-term marine biodiversity observation. Analyzing structural face categories increases sensitivity and reproducibility while maintaining manageable effort, providing a practical design for regional and global observatories. These methodological refinements align with emerging observatory networks such as ARMS-MBON and OBON, promoting interoperable and standardized biodiversity monitoring across space and time.

## Author contributions

AC and VD conceived and designed the study. AC, AH, DG and FZ chose the sampling sites. AC, AH, CM, DG, EB, Fl.M, FZ, LV, LP, MS, PM, SC, SR, TL, VC and VR performed the fieldwork (scuba-diving) and/or ARMS processing (fractioning, scraping, fixation, and photography). VD conceived and developed the metabarcoding protocol. Fl.M, Fa.M and CC did the molecular work. EM did the bioinformatics. JMGO provided environmental variables for all sites. AC, EB, DG, ML and SR developed the protocol and/or contributed to the analyses of photography. LP trained the consortium members on ARMS processing. AC performed all statistical analyses and figures. AC and VD wrote the original draft. EB, EM, JMGO, PM, TL, SR and VR contributed to further writing and editing. All authors approved the final version of the manuscript.

## Conflict of interest

The authors declare no conflict of interest

## Acknowledgements

We thank Thierry Thibaut, Jean-Georges Harmelin, and Jean Vacelet for their assistance with photographic analyses and Federica Costantini and Marco Abbiatti for help in ARMS dismantlement in Marseille during a training organized by IMBE and OSU Pythéas. Molecular data used in this study were produced by the molecular facilities of CIRAD (Montferrier-sur-Lez) and BMC (IMBE, Marseille). Scuba diving were OSU Pythéas. This study has been conducted using E.U. Copernicus Marine Service Information of product MEDSEA_MULTIYEAR_PHY_006_004 and MEDSEA_ANALYSIS_FORECAST_BIO_006_008; https://doi.org/10.25423/CMCC/MEDSEA_MULTIYEAR_PHY_006_004_E3R1, ; https://doi.org/10.25423/cmcc/medsea_multiyear_bgc_006_008_medbfm3

## Data availability statement

Sequence data have been deposited in the NCBI Sequence Read Archive (SRA) under the BioProject ID PRJNA1231031.

---

**Figure S1:**
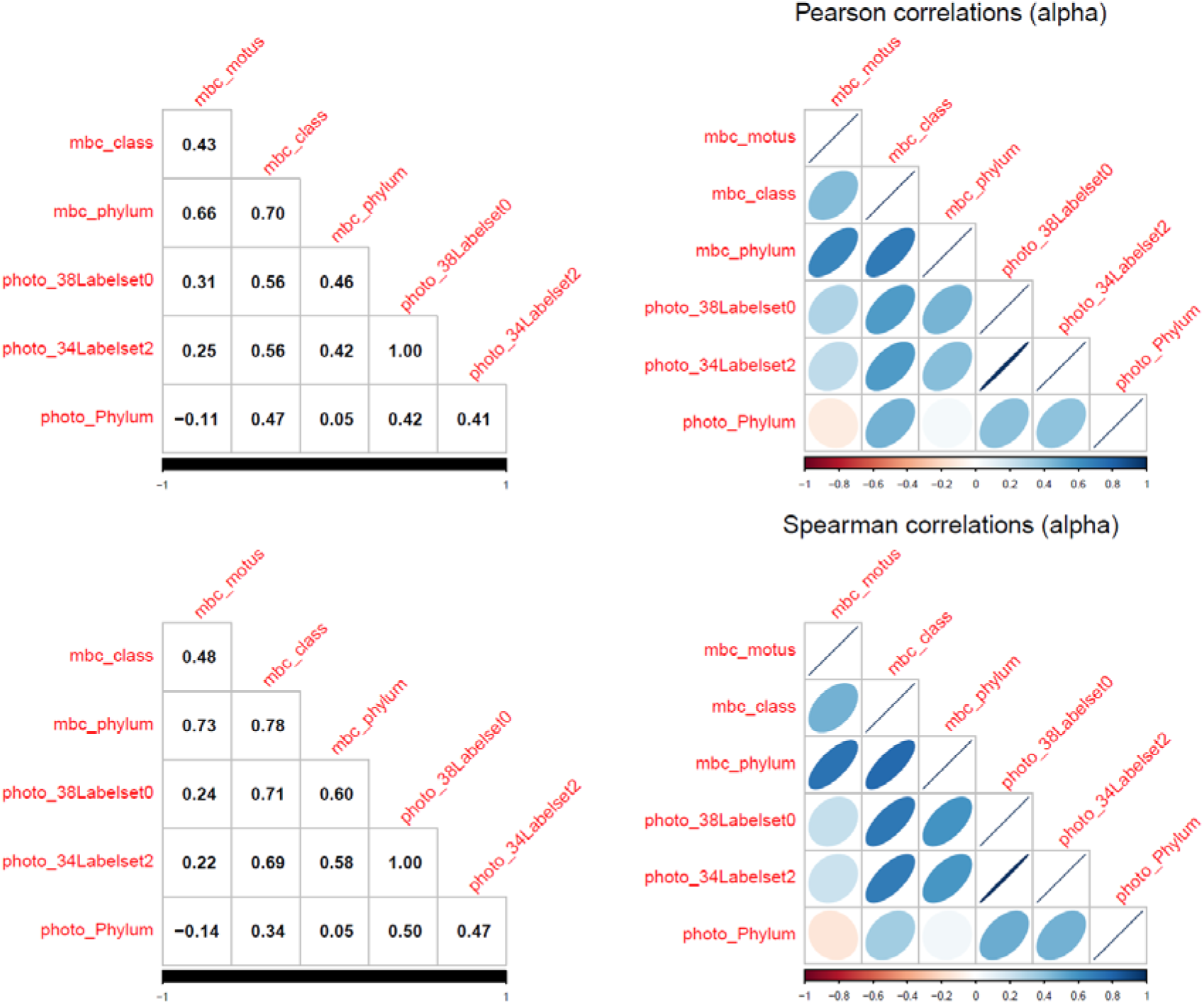
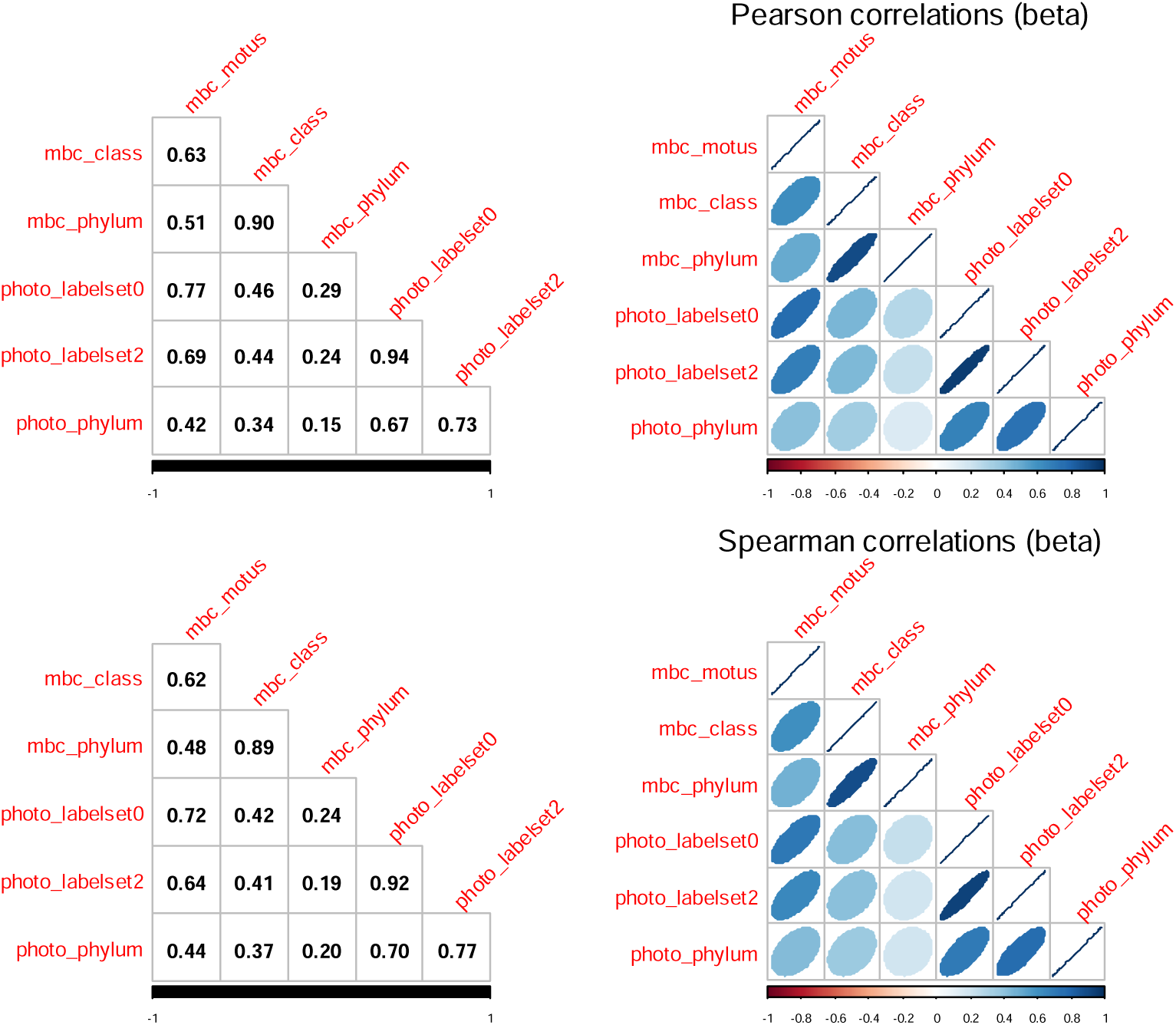
Correlation plots of α-and β-diversity among photographic and metabarcoding datasets at the 10 sites level. Pairwise correlations of α-diversity (upper matrices) and β-diversity (lower matrices) between datasets defined by three taxonomic-rank levels for both photographic and metabarcoding data, drawn by the R package corrplot (version 0.95). Correlations on Spearman’s rank coefficients were similar to Pearson’s correlations. For α-diversity, only matrix cells with correlation coefficients ≥ 0.69 are significant at p < 0.05, indicating that only metabarcoding data aggregated at the class level are positively correlated with photographic data for site richness. For β-diversity, the threshold is 0.294. The metabarcoding dataset used here corresponds to the 17-faces design (for comparisons with photography).

**Table S1:** Study sites and sampling details for 2019. For each site, the table reports depth, geographical coordinates (decimal latitude and longitude), dates of immersion and emersion. In each site in 2019, three ARMS were analysedanalyzed the same way: for the sessile fraction of each ARMS, faces were scraped and grouped according to their orientation and compartmentation into five categories (see the methods), then five DNA extractions and metabarcoding were carried out as explained in the methods. Two sites (BAN1 and VIL3) are reported here for information on the SEAMoBB project, but were not included in analyses presented in this study, to ensure comparability with photography and separate face metabarcoding.

| Site code | Site name | Region | Latitude | Longitude | Depth | Date in | Date out |
| --- | --- | --- | --- | --- | --- | --- | --- |
| BAN1 | Port-vendres S | Banyuls-sur-Mer | 42,51327 | 3,13743 | 17 m | 17/04/2018 | 20/02/2019 |
| BAN2 | Port-vendres N | Banyuls-sur-Mer | 42,52518 | 3,11050 | 17 m | 12/04/2018 | 19/02/2019 |
| BAN3 | Cerbère N | Banyuls-sur-Mer | 42,44818 | 3,17212 | 17 m | 19/03/2018 | 20/02/2019 |
| BAN4 | Cerbère S | Banyuls-sur-Mer | 42,44262 | 3,17405 | 17 m | 19/03/2018 | 20/02/2019 |
| MAR1 | Veyron | Marseille | 43,20683 | 5,25350 | 17,5 m | 07/05/2018 | 19/04/2019 |
| MAR2 | Digue des catalans | Marseille | 43,29167 | 5,34558 | 19 m | 05/03/2018 | 26/03/2019 |
| MAR3 | Niolon | Marseille | 43,34150 | 5,26525 | 16,5 m | 09/05/2018 | 26/03/2019 |
| MAR4 | Plane | Marseille | 43,19067 | 5,38233 | 17,5 m | 05/03/2018 | 19/04/2019 |
| VIL1 | Pointe de la gavinette | Villefranche-sur-Mer | 43,68683 | 7,32138 | 20 m | 13/03/2018 | 04/03/2019 |
| VIL2 | Phare du cap ferrat | Villefranche-sur-Mer | 43,67427 | 7,32803 | 18 m | 15/03/2018 | 06/03/2019 |
| VIL3 | Grotte à corail | Villefranche-sur-Mer | 43,68945 | 7,30883 | 17 m | 13/03/2018 | 05/03/2019 |
| VIL4 | Cap ferrat est | Villefranche-sur-Mer | 43,67675 | 7,33508 | 18 m | 14/03/2018 | 06/03/2019 |

## Notes

**Funding:** This work was supported by the MONITOBEN and MONITOBEN-2 projects, funded by EMBRC-France (OOB_EMBRC FR_AAP2018_n°2179), and by the SEAMoBB project (Solutions for sEmi-Automated Monitoring of Benthic Biodiversity), co-funded by ERA-Net Mar-TERA (id. 145), the French National Research Agency (ANR; Grants ANR-17-MART0001-01, ANR-17-MART0001-02 and ANR-17-MART0001-03), and the Centro para el Desarrollo Tecnológico e Industrial (CDTI, Spain; Project SERA-20181031 “Soluciones para el Monitoreo Semi-Automático de la Biodiversidad Bentónica”). V.R. acknowledge partial financial support from the European Space Agency (ESA Contract No.4000141547/23/I-DT) through the 4DMED-Sea project. The project leading to this publication has received funding from European FEDER Fund under project 1166-39417.

https://www.ncbi.nlm.nih.gov/bioproject/1231031

### Competing Interest Statement

The authors have declared no competing interest.

